# Historical contingency drives compensatory evolution and rare reversal of phage resistance

**DOI:** 10.1101/2022.02.10.479996

**Authors:** Reena Debray, Nina De Luna, Britt Koskella

**Author notes:** Corresponding author*: Reena Debray, Mailing address: 3040 Valley Life Sciences Building #3140, Berkeley, CA, 94720. Contributing authors*: Nina De Luna, Britt Koskella.

## Abstract

Bacteria and lytic viruses (phages) engage in highly dynamic coevolutionary interactions over time, yet we have little idea of how transient selection by phages might shape the future evolutionary trajectories of their host populations. To explore this question, we generated genetically diverse phage-resistant mutants of the bacterium *Pseudomonas syringae*. We subjected the panel of mutants to prolonged experimental evolution in the absence of phages. Some populations re-evolved phage sensitivity, while others acquired compensatory mutations that reduced the costs of resistance without altering resistance levels. To ask whether these outcomes were driven by the initial genetic mechanisms of resistance, we next evolved independent replicates of each mutant lineage in the absence of phages. We found a strong signature of historical contingency: some mutations were highly reversible across replicate populations, while others were highly entrenched. Through whole-genome sequencing of bacteria over time, we also found that populations with resistance mutations in the same gene acquired more parallel sets of mutations than populations with resistance mutations in different genes, suggesting that compensatory adaptation is also contingent on how resistance initially evolved. Our study identifies an evolutionary ratchet in bacteria-phage coevolution, and may explain previous observations that resistance persists over time in some bacterial populations but is lost in others. We add to a growing body of work describing the key role of phages in the ecological and evolutionary dynamics of their host communities. Beyond this specific trait, our study provides new insight into the genetic architecture of historical contingency, a crucial component of interpreting and predicting evolution.

## INTRODUCTION

Pathogens are ubiquitous and exert strong selection on their hosts to evade infection (Stenseth and Maynard Smith 1984; Gómez, Verdú, and Perfectti 2010). These selection pressures are constantly in flux, and defense-related traits are often detrimental when pathogens are not present at high levels (Clay and Kover 1996; Sheldon and Verhulst 1996). The loss of costly resistance under relaxed selection has been the focus of a plethora of theoretical and empirical studies, in large part because it helps to explain the observed coexistence of resistant and sensitive host types in many natural populations (Waterbury and Valois 1993; Stahl et al. 1999; Rodriguez-Brito et al. 2010; Lourenço et al. 2020). Whether host populations readily regress to susceptibility after escape from pathogen pressure or retain a signature of their coevolutionary history will depend on several factors, including the environmental conditions and the strength of selection. For example, resistance may persist if it is not costly to maintain; or if compensatory mutations reduce the fitness costs without reversing the trait itself, as is often observed in drug-resistant bacteria (Faria et al. 2015; Durão, Balbontín, and Gordo 2018). In other cases, even when reversion to sensitivity would be favorable, it may not be possible if the host population has since acquired other mutations that would be deleterious on the wild-type genetic background (Shah, McCandlish, and Plotkin 2015).

In a more general sense, any mutation that affects an organism’s phenotype and is selected for (even temporarily) can alter the selection acting on subsequent mutations, and can therefore shape the evolutionary trajectory of the population. For example, garter snakes that prey on tetrodotoxin-bearing newts have evolved high levels of toxin resistance, but only within lineages that already carried a prior substitution – an ancient modification to a sodium channel that took place long before the newts arose (McGlothlin et al. 2016). In a laboratory evolution experiment with *Escherichia coli*, different substitutions in a DNA topoisomerase enzyme were shown to have different consequences for the subsequent accumulation of other beneficial mutations. In fact, this second-order selection for evolvability was more important than the initial effects of the substitutions themselves in determining which lineages prevailed in the long term (Woods et al. 2011). Historical contingencies such as these can make it particularly challenging to interpret patterns of genetic divergence across populations or predict the lasting consequences of short-term coevolutionary interactions.

To explore how previous selection by phages can alter future bacterial evolution, we tracked phage resistance over time in experimental evolution populations of the bacterium *Pseudomonas syringae*. Bacteria and phages are a tractable model system frequently used for studying coevolution in the laboratory (Brockhurst et al. 2007; Dennehy 2012). Phages initiate infection by recognizing and binding to proteins on the surfaces of bacterial cells. Bacteria can evade phages by altering or deleting phage receptors, but in doing so, often compromise other fitness-related traits such as nutrient uptake, adhesion, and virulence (Dy et al. 2014; Mangalea and Duerkop 2020). A few studies have propagated resistant bacteria for many generations in the absence of phages, but whether phage sensitivity re-emerges has remained inconsistent and difficult to explain (Meyer et al. 2010; Avrani and Lindell 2015). For example, over 45,000 generations of relaxed selection did not reduce the resistance of *Escherichia coli* to T6 phage (Meyer et al. 2010). In contrast, some experimental populations of *Prochlorococcus* became less resistant in the absence of phage, but these changes were difficult to explain in terms of compensatory adaptation, as they often occurred independently of fitness gains (Avrani and Lindell 2015).

On the basis of these observations, we hypothesized that bacteria can access many different genetic pathways to phage resistance, each with different implications for the subsequent evolutionary potential of the bacteria, including whether they compensate for fitness costs by re-evolving phage sensitivity. To test this idea, we isolated and sequenced *P. syringae* colonies that had evolved resistance across a panel of lytic phages and measured the fitness costs of each resistance mutation. We then used each resistant strain to seed a different population that was experimentally evolved in the absence of phages. Through a series of laboratory evolution experiments, we demonstrate that phage resistance can either be reversible or entrenched depending on the initial genetic path to resistance.

## RESULTS

### Single-step selection for phage resistance

To explore whether the long-term outcomes of phage resistance were contingent on the genetic underpinnings of their initial resistance (and/or the fitness costs associated with resistance), we first sought to generate a panel of resistant bacterial strains with substantial variation both in resistance mechanisms and in fitness. Using five different lytic phages (see Methods for details), we selected 22 phage-resistant colonies of *Pseudomonas syringae* pv. tomato DC3000 through soft agar overlays. Whole-genome sequencing of these 22 resistant strains revealed 17 unique resistance mutations (positions 1184510 and 5662024 each appeared three times in the panel, and one colony had no detectable fixed genetic differences from the ancestor despite apparent phenotypic resistance).

In contrast to the diversity of exact mutations, there was relatively high convergence in the genes in which they were found. The mutations in this study were all located in one of four genes, all involved in biosynthesis of outer membrane lipopolysaccharide molecules (Figure **1a**). Resistance mutations included single-nucleotide missense mutations, single-nucleotide nonsense mutations, and insertions and deletions of varying sizes. When resistant mutants were tested in the absence of phages, these strains grew more slowly than their phage-sensitive ancestor, indicating trade-offs between resistance and other aspects of fitness (one-sample t-test, t = −8.405, *p* <0.001, Figure **1b**). Variation in growth rates could not be explained by the genes in which those mutations occurred (ANOVA, F = 0.673, df = 3, *p* = 0.580), indicating that different resistance mutations in the same gene could have different impacts on fitness. This lack of systematic differences in fitness costs across genes underlying resistance allowed us to tease apart the effects of underlying genetic mechanisms and fitness costs on the maintenance of phage resistance, as these two predictors were not confounded with each other.

**Figure 1.**
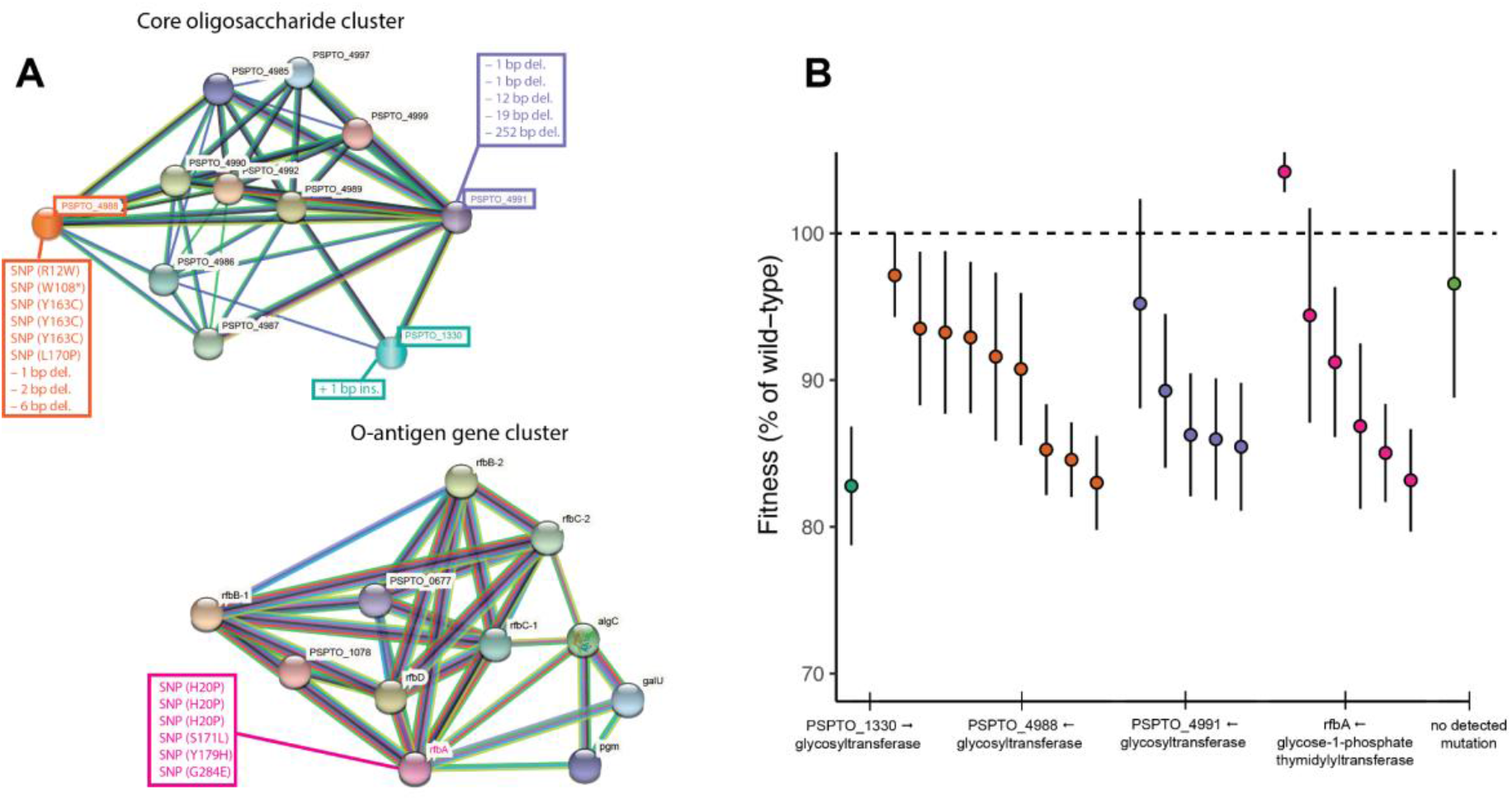
Genetic mechanisms and fitness costs of phage resistance. **(a)** Bacterial proteins implicated in phage resistance in this study and their predicted functional partners according to the Search Tool for the Retrieval of Interacting Genes/Proteins (STRING) database. Colored text boxes list the specific mutations observed in this study (n=21). In the case of SNPs, the location and the nature of the mutation is indicated in parentheses; e.g. R12W indicates a change from arginine to tryptophan at the twelfth position, and W108* indicates a change from tryptophan to a stop codon. **(b)** Population growth rates of resistant strains in the absence of phage, based on a logistic model fitted to a 40-hour growth curve (n=22 strains). Error bars represent variation across technical replicates. Bacterial colonies with resistance mutations in the same gene are denoted by the color scheme (which corresponds to the proteins in panel a), and the arrow next to each gene name indicates orientation of the gene (positive or negative strand).

### Multiple evolutionary paths to recover fitness costs

To test whether phage sensitivity tends to re-evolve in the absence of phages as a result of these fitness costs, we inoculated 22 experimental microcosms of King’s B medium each with one of the resistant colonies. We also inoculated 6 microcosms with the phage-sensitive ancestor to assess changes due to overall adaptation to the environment over the course of experimental evolution, for a total of 28 experimentally evolving populations. Populations were transferred to fresh media every 3 days for a total of 36 days. Over the course of the experiment, the difference in growth rates between the populations initiated with phage-resistant bacteria and those initiated with phage-sensitive (ancestral) bacteria gradually narrowed, and by the end of the experiment there were no detectable costs of resistance observed (Figure **2a**). Changes in fitness followed a diminishing returns pattern of adaptation: populations with the lowest initial fitness made the greatest fitness gains over time (ANOVA, F = 33.226, *p* < 0.001; Figure **S1**).

**Figure 2.**
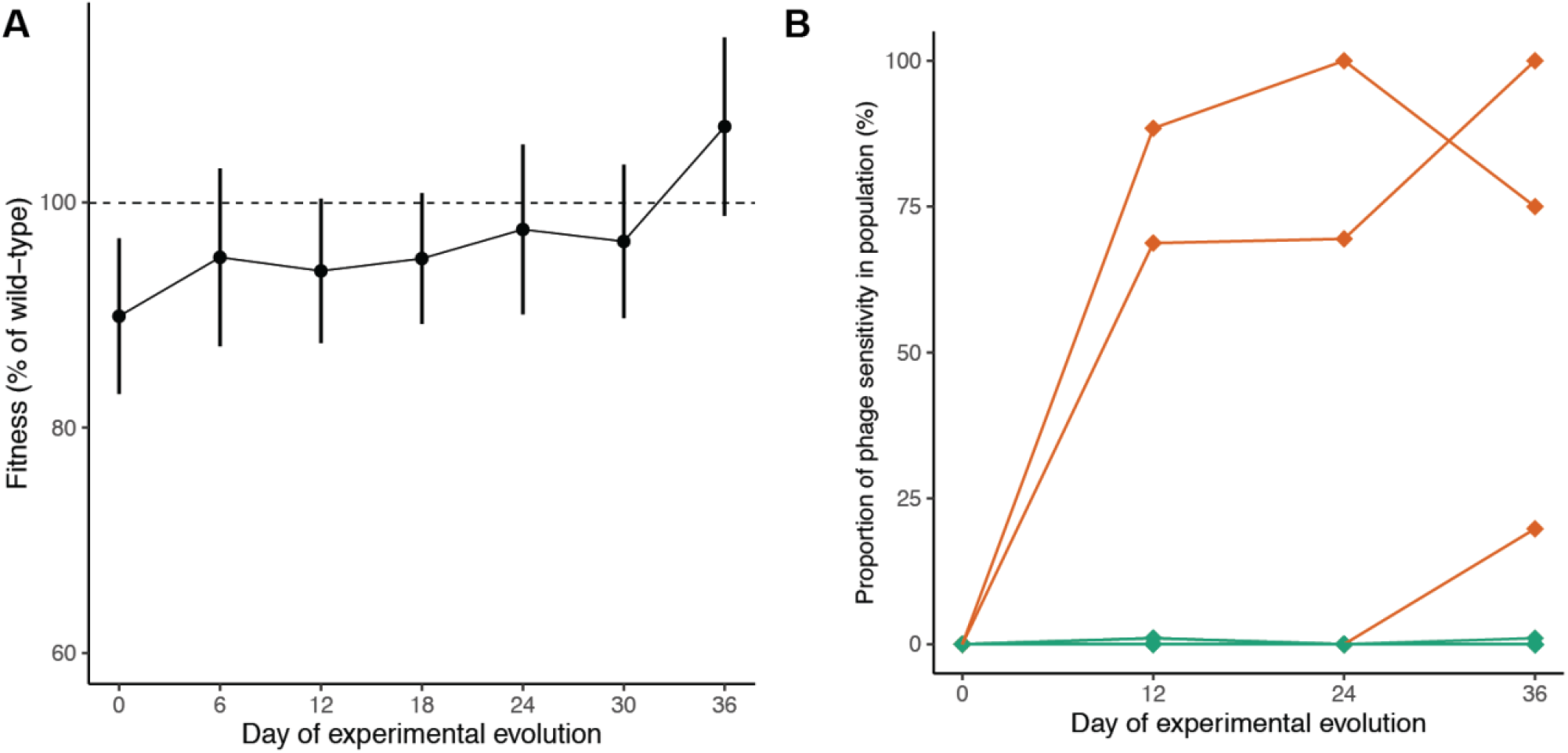
Phenotypic changes in bacterial populations during experimental evolution in the absence of phages. **(a)** Population growth rates of populations inoculated with phage-resistant colonies over experimental evolution in the absence of phages, calculated based on optical density measurements in a microplate reader over 40 hours of growth (n=22). Points indicate mean fitness among populations sampled at the same day and error bars indicate standard deviation. Values are normalized to the growth rates of the phage-sensitive control populations sampled at the same point in experimental evolution. **(b)** Proportion of colonies in each population that were scored as phage-sensitive over time, as indicated by disruption in bacterial growth upon encountering phage on an agar plate (n=22 populations with 96 colonies sampled per population). Orange lines indicate populations from which sensitive colonies were isolated and verified in a microplate assay.

Phage-sensitive phenotypes re-evolved under relaxed selection, but only rarely. A total of 5 populations (out of 22 initially resistant populations) contained colonies whose growth appeared to be disrupted by phage on agar plates. Of these, only 3 populations could be further validated for phage sensitivity based on measurements of population growth over time in the presence and absence of phages (Figure **2b**). Interestingly, the overall increase in bacterial fitness over time could not be attributed to the re-evolution of phage sensitivity alone. While there was a main effect of the day of experimental evolution on bacterial fitness, the rate of change did not differ among populations that re-evolved sensitivity and those that remained resistant (ANOVA, F = 2.146 for the interaction term, *p* = 0.145). This observation suggested that the resistant populations may be able to access compensatory mutations that lessen the costs of phage resistance without reversing it.

In cases where phage sensitivity re-evolved within initially resistant populations, we isolated and sequenced the genomes of phage-sensitive colonies to determine the mechanism of reversion. The population “SNK7”, which originally acquired resistance through a single base-pair deletion in the glycosyltransferase-encoding gene PSPTO_4991, later acquired a single base-pair insertion 5 bases away that restored the reading frame. In contrast, the other two populations acquired mutations that fell outside of their initial resistance genes. VCM4, which had become phage-resistant through a substitution in the *rfbA* gene, evolved additional mutations in two other membrane transport proteins. QAC5, which had evolved resistance through a single base-pair insertion in the glycosyltransferase-encoding gene PSPTO_1330, acquired a mutation in a membrane transport protein as well as several intergenic mutations whose function (if any) is unknown.

We had predicted that populations with particularly costly resistance would be most likely to re-evolve sensitivity due to strong selection. However, we did not observe a relationship between initial costs of resistance and the evolution of phage sensitivity (Welch’s unequal variances t-test, t = 1.818, *p* = 0.138). We also asked whether phage sensitivity would be more likely to re-emerge among populations with particular resistance genes. While the unequal distribution of phage resistance outcomes makes it difficult to entirely rule out this possibility, the three populations that re-evolved sensitivity in this study did so from three different initial resistance genes, providing no evidence that trait reversion is historically contingent at the gene level (Table **1**). However, we noticed two important observations that warranted further study.

**Table 1.** Genetic mechanisms and evolutionary outcomes of phage-resistant populations.

| **Population** | **Resistance gene** | **Position of resistance mutation** | **Type of resistance mutation** | **Outcome in original evolution experiment** |
| --- | --- | --- | --- | --- |
| FMS6 | *rfbA* | 1183718 | SNP | Remained resistant |
| FMS4 | *rfbA* | 1184034 | SNP | Remained resistant |
| VCM4 | *rfbA* | 1184057 | SNP | Re-evolved sensitivity |
| MS10 | *rfbA* | 1184510 | SNP | Remained resistant |
| MS2 | *rfbA* | 1184510 | SNP | Remained resistant |
| QAC4 | *rfbA* | 1184510 | SNP | Remained resistant |
| QAC5 | PSPTO_1330 | 1461031 | 1 bp ins. | Re-evolved sensitivity |
| VCM19 | PSPTO_4988 | 5661617 | 2 bp del. | Remained resistant |
| MS12 | PSPTO_4988 | 5661621 | 1 bp del. | Remained resistant |
| FMS13 | PSPTO_4988 | 5661925 | 6 bp del. | Remained resistant |
| SNK12 | PSPTO_4988 | 5662003 | SNP | Remained resistant |
| QAC3 | PSPTO_4988 | 5662024 | SNP | Remained resistant |
| SNK11 | PSPTO_4988 | 5662024 | SNP | Remained resistant |
| VCM17 | PSPTO_4988 | 5662024 | SNP | Remained resistant |
| SNK6 | PSPTO_4988 | 5662188 | SNP | Remained resistant |
| MS1 | PSPTO_4988 | 5662478 | SNP | Remained resistant |
| FMS12 | PSPTO_4991 | 5664934 | 19 bp del. | Remained resistant |
| FMS9 | PSPTO_4991 | 5665124 | 252 bp del. | Remained resistant |
| SNK7 | PSPTO_4991 | 5665501 | 1 bp del. | Re-evolved sensitivity |
| FMS11 | PSPTO_4991 | 5665743 | 1 bp del. | Remained resistant |
| FMS16 | PSPTO_4991 | 5665890 | 12 bp del. | Remained resistant |

First, our study included four populations that had originally acquired resistance through large deletions in oligosaccharide synthesis genes, spanning from 6 base pairs to 252 base pairs in length. None of these populations re-evolved sensitivity, whether by re-insertion of the missing sequence or through any other means. Second, by chance our study included two sets of three populations each with identical resistance mutations. In both cases, these genetic triplets all followed the same evolutionary outcome (remaining resistant). Given that re-evolving phage sensitivity was generally rare in this study, it was not apparent whether these two observations were simply probabilistic, or whether they pointed to historical contingency at the level of the individual mutation rather than the resistance gene as a whole. As this experiment was not originally set up to test this possibility, we developed an experimental design that would test the effects of exact resistance mutations on phage resistance outcomes.

### Replay of experimental evolution

The initial evolution experiment revealed that the identity of the gene conferring phage resistance did not explain whether bacteria remained resistant or re-evolved sensitivity. To ask whether phage resistance outcomes were contingent on the exact resistance mutation instead, we returned to the progenitor frozen stocks of the 3 resistant strains that had re-evolved sensitivity during experimental evolution. We also included 3 resistant strains that were never observed to re-evolve sensitivity, selecting the closest possible genetic matches to the strains that did. For example, VCM4 and MS2 both originally acquired resistance through point mutations in the *rfbA* gene, yet VCM4 re-evolved phage sensitivity while MS2 did not.

We used each of these 6 resistant mutants (“founders”) to seed 10 replicate populations, resulting in a total of 60 experimentally evolving populations. As above, these populations were maintained in experimental microcosms of King’s B medium and transferred every 3 days for 36 days. At the end of the replay experiment, founder identity accounted for the majority of variance among populations in the prevalence of phage sensitivity, with evolutionary stochasticity playing a smaller role. In fact, populations derived from a founder that had re-evolved sensitivity in the original experiment were far more likely to re-evolve sensitivity in the replay experiment as well (Welch’s unequal variances t-test, t = 4.744, *p* < 0.001; Figure **3**).

**Figure 3.**
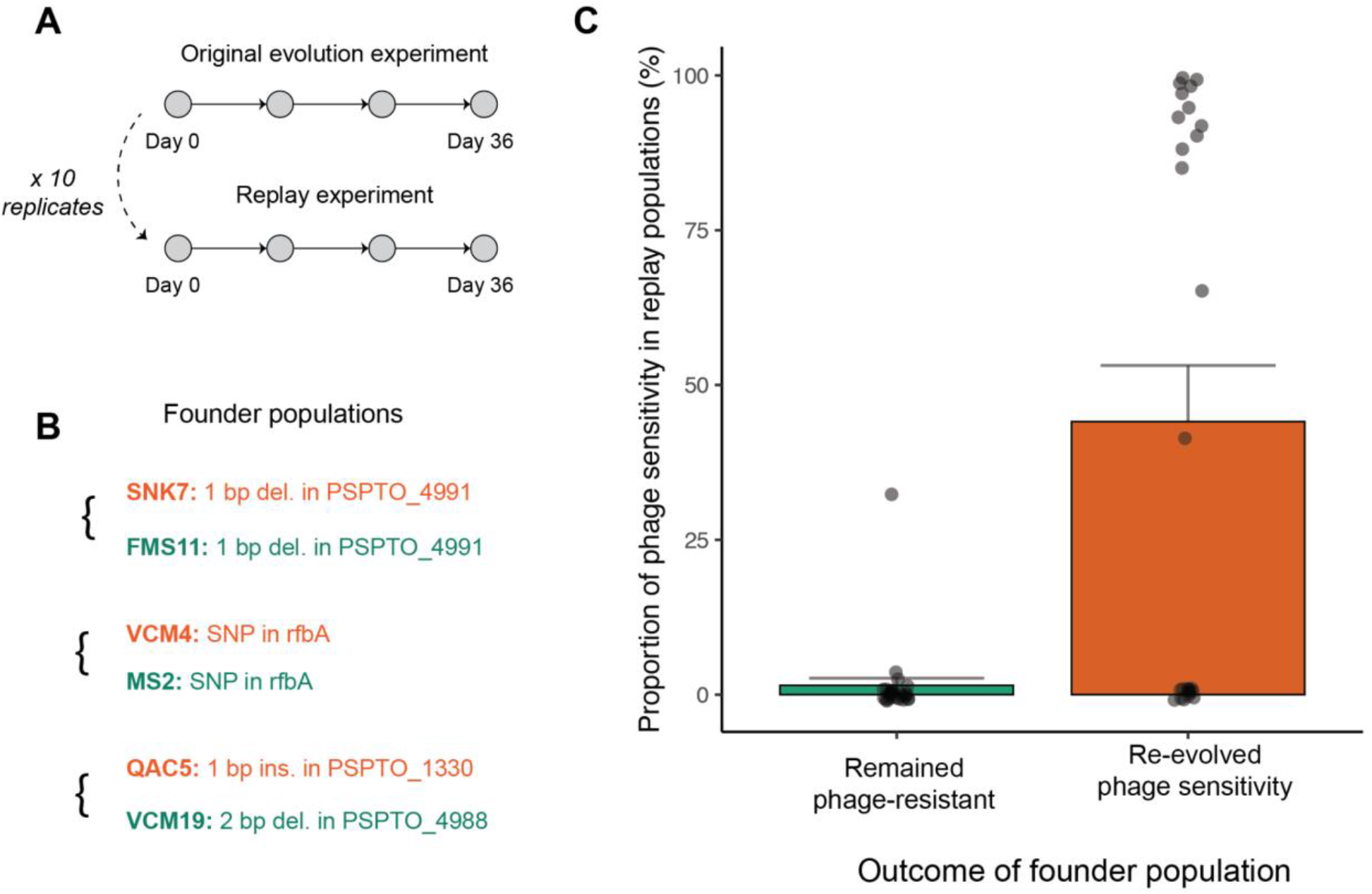
Phage resistance outcomes in the replay experiment mirror the outcomes of their founding population. **(a)** Populations in the replay experiment were founded by frozen stocks of resistant colonies prior to their evolution in the absence of phages. Each founder was used to generate 10 replicate populations from genetically identical starting points. **(b)** Resistance mutations of the 6 founders. Each colony that had re-evolved sensitivity in the original experiment (orange) was paired with a genetically similar colony that had remained resistant (green). Of note, QAC5 was the only colony in the study with a mutation in PSPTO_1330, so it was matched with a population that also had a small frameshift mutation, but in a different glycosyltransferase-encoding gene. **(c)**Proportion of colonies in each population that were scored as phage-sensitive after 36 days in the absence of phages, as indicated by disruption in bacterial growth upon encountering phage on an agar plate (n=54 populations with 96 colonies sampled per population).

### Historical contingency in genome evolution

To characterize genetic changes during experimental evolution, we deep-sequenced all of the populations of the original evolution experiment at two time-points: early in experimental evolution (Day 6) and at the end of the experiment (Day 36). The evolving populations fixed relatively few mutations in protein-coding genes by the end of the experiment (mean = 0.71 ± 0.66 s.d.), but acquired and maintained polymorphisms at many loci (mean = 22.61 ± 7.20 s.d.). Across all populations, the most frequent mutations occurred in effector proteins, membrane-bound enzymes, and transcription factors (Table **S1**). Additionally, several populations acquired substitutions in the DNA mismatch repair proteins *mutS* or *mutL* during experimental evolution. While none of these mutations fixed within their respective populations, populations with a mutation in *mutS* or *mutL* acquired more fixed mutations or polymorphisms overall than the other populations by the end of the experiment (Student’s t-test, t = 2.746, *p* = 0.005), suggesting that they may contain ‘hypermutator’ lineages with elevated mutation rates (Shaver et al. 2002; Harris, Flynn, and Cooper 2021).

To characterize the molecular basis of compensatory evolution in phage-resistant bacteria, we first considered that compensatory mutations in other systems are often related to biochemical protein stability (Bridgham, Ortlund, and Thornton 2009; Gong, Suchard, and Bloom 2013). We therefore predicted that different mechanisms of resistance would likely require different compensatory mutations to ameliorate their costs. We would then expect populations with the same resistance gene to exhibit greater evolutionary parallelism than populations with different resistance genes. We calculated the Sørenson-Dice similarity coefficient, which describes the proportion of mutated genes that two populations have in common, controlling for their initial genetic backgrounds. At day 6 of the experiment, populations with the same resistance mechanisms had acquired more similar sets of mutations than populations with different resistance mechanisms (randomization test, *p* < 0.001, Figure **4a**). For example, populations that had originally acquired phage resistance through a mutation in the glycosyltransferase-encoding enzyme PSPTO_4991 evolved particularly similarly to one another, with many genes acquiring mutations in either all or none of the populations (see Figure **4b**for a diagram of genetic differences between evolved and ancestral populations organized by initial resistance gene). After an additional 30 days of experimental evolution in the same environment, there was no remaining signature of the initial resistance gene on pairwise similarity coefficients (randomization test, *p* = 0.710, Figure **S2**).

**Figure 4.**
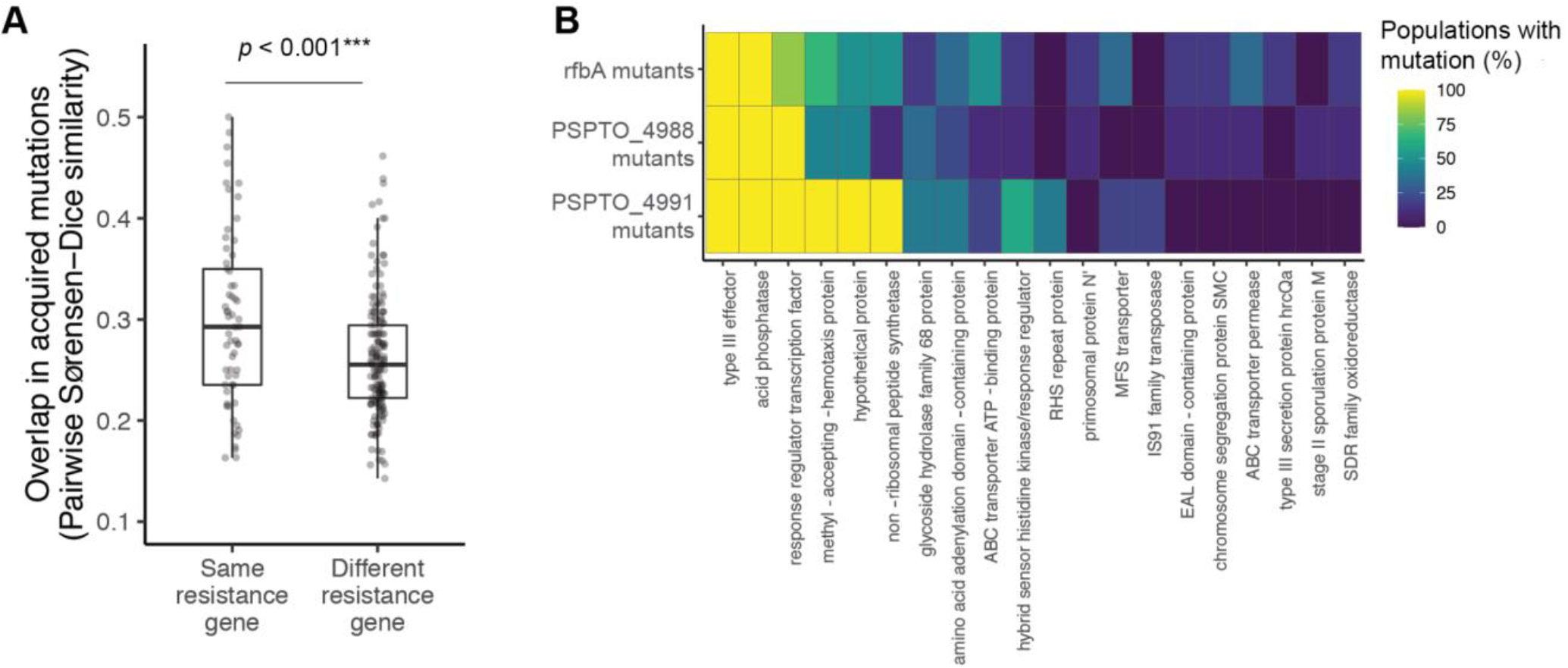
Genetic contingency in identity of mutated genes during experimental evolution. **(a)** Pairwise similarity coefficients among pairs of evolved populations at day 6 of experimental evolution (n=22). Statistical significance was assessed by randomizing whether pairs were labeled as having the same or different resistance genes and recalculating their similarity coefficients for 10,000 permutations. **(b)** Heatmap depicting the relationship between initial resistance genes and mutations acquired during experimental evolution (n=20). Rows represent all populations with resistance mutations in the same gene (note that resistance genes represented by fewer than 2 populations are not pictured, as there was no way to assess parallelism in these cases). Columns represent the top 20 genes that were most frequently mutated across populations by day 6 of experimental evolution. Colors indicate the percentage of populations of each resistance gene that had acquired one or more mutations by day 6 of experimental evolution.

## DISCUSSION

Phages play a central role in shaping bacterial evolution in nature, with critical implications for bacteria-bacteria and bacteria-host interactions (Thurber 2009; Koskella and Brockhurst 2014). Predation by phages can maintain population and community diversity in their bacterial hosts (Weinbauer and Rassoulzadegan 2004), regulate the dissemination of antibiotic resistance genes (Burmeister et al. 2020; LeGault et al. 2021), select for hypermutator strains (Pal et al. 2007), and alter competitive outcomes among bacterial species (Bohannan and Lenski 2000). Here, we show that even transient exposure to phages can have lasting consequences for the evolutionary trajectories of bacterial populations. Through a series of laboratory evolution experiments, we demonstrate that phage resistance in the plant pathogenic bacterium *Pseudomonas syringae* can be reversible or entrenched depending on the original genetic path to resistance.

### Mechanisms and costs of phage resistance

Phage resistance in our study primarily occurred through mutations in lipopolysaccharide biosynthesis genes, which have been previously implicated in phage resistance in this bacterial species and others (Picken and Beacham 1977; Evans et al. 2010; Meaden, Paszkiewicz, and Koskella 2015; Kulikov et al. 2019). One population did not appear to have any fixed genetic differences from the ancestor despite its resistant phenotype. Resistance in this case may be conferred by mutations that are not adequately resolved by short-read sequencing, such as copy number variation or sequence region inversions, or through unstable genetic changes such as phase variations (Kircher and Kelso 2010; Bull et al. 2014). Of note, this population remained phenotypically resistant throughout the evolution experiment, possibly because it did not experience detectable fitness costs relative to its phage-sensitive ancestor.

Even though the resistance mutations in our study were concentrated within a handful of genes, they occurred at many unique positions, and altered the amino acid sequences in a variety of ways, including missense and nonsense substitutions and frameshift mutations of vastly different sizes. Convergence at the gene level and diversity at the sequence level is consistent with the expectation that phage infection relies on specific receptor structures, and that recognition can easily be disrupted by modifying or deleting these structures in one of many ways (Dy et al. 2014). However, resistance was generally costly in the absence of phages, as is often observed in this system and others (Koskella et al. 2012; Meaden, Paszkiewicz, and Koskella 2015; Scanlan, Buckling, and Hall 2015; Burmeister, Sullivan, and Lenski 2020; Burmeister et al. 2020). Such costs of resistance may stem from alterations to lipopolysaccharide molecules that destabilize the bacterial membrane or reduce surface adhesion (Mangalea and Duerkop 2020). These fitness costs suggest that resistance will be selected against in the absence of phage pressure, yet previous studies addressing this question have produced inconsistent results (Meyer et al. 2010; Avrani and Lindell 2015; Wielgoss et al. 2016). When we propagated the bacterial populations for an extended period in the absence of phages, we observed that phage sensitivity re-appeared and swept to high frequencies in several populations, but that the majority of populations remained resistant. In the case of one population, this reversion to sensitivity was due to a mutation that restored the reading frame of the original sequence, but in other cases, populations re-evolved phage sensitivity without reversing the original mutation.

### Historical contingency in phenotypic evolution

The diversity of phage-resistant mutants in this study allowed us to ask whether the re-evolution of phage sensitivity was contingent on the mechanisms and/or costs of phage resistance. Costs of trait maintenance are often expected to predict trait loss under relaxed selection (Lahti et al. 2009; Meyer et al. 2010), yet we did not observe a relationship between the magnitude of fitness costs and the re-evolution of phage sensitivity in our study. This observation, along with the fact that populations that did not re-evolve sensitivity nevertheless improved their fitness to match their phage-sensitive counterparts, suggests the existence of compensatory mutations that reduce the costs of resistance (Pennings, Brandon Ogbunugafor, and Hershberg, n.d.). Therefore, it appears that trade-offs between phage resistance and other aspects of fitness can be strong, yet bacteria are able to access two evolutionary pathways (reversion and compensation) in response.

We hypothesized that the probabilities of these two pathways could be contingent on the genetic mechanism underlying phage resistance. Some traits might have a greater supply of reversion mutations than others; for example, there may be more mutations that restore the ancestral expression levels of a gene than those that reconstitute the exact three-dimensional structure of a receptor (Wielgoss et al. 2016). Similarly, resistance mechanisms may impact bacterial fitness in different ways, thus requiring different sets of compensatory mutations to restore their costs (Rojas Echenique et al. 2019). We did not find an overall correlation between the initial resistance gene and whether phage sensitivity re-evolved, but with two important caveats. First, when resistance was acquired through a large deletion in a receptor biosynthesis gene, it was never reversed, whether through a complementary insertion or other mutations. This suggests that the mutations that would reverse such a large change are so vanishingly rare that other compensatory mutations are much more likely to appear first. Compensatory mutations could eventually restore fitness levels to a point where phage sensitivity is no longer advantageous (Teotónio and Rose 2007; McCandlish, Shah, and Plotkin 2016; Pennings, Brandon Ogbunugafor, and Hershberg, n.d.), or may even be disadvantageous or lethal on the ancestral background, further discouraging reversion of the original trait (Rojas Echenique et al. 2019).

Our second observation was that the genetic replicates in the mutant panel (i.e. the populations that independently acquired resistance via the same mutation) all followed the same phenotypic trajectories. Specifically, they all remained resistant throughout experimental evolution in the absence of phage. This suggested that historical contingencies – if they existed here – might be generated not from the identity of the resistance gene, but from the exact genetic sequence. Even within the same resistance gene, some mutations might be more reversible than others. As the original experiment was not explicitly focused on this possibility, we designed a follow-up experiment that “replayed” experimental evolution 10 times per founding genotype (inspired by Stephen Jay Gould’s famous thought experiment about replaying the tape of life, (Gould 1990)). Strikingly, we found that the evolutionary outcomes of the populations in our replay experiment closely mirrored those of their founders. Populations whose founders had re-evolved phage sensitivity also tended to re-evolve phage sensitivity at high rates, while populations whose founders had remained resistant tended to remain resistant as well.

### Historical contingency in molecular evolution

Variation in the reversibility of different resistance mutations may occur if different sets of compensatory mutations are required to restore their costs. To explore this possibility, we compared the similarity of acquired mutations among pairs of the experimentally evolving populations. We found that populations with the same initial resistance gene evolved more in parallel with one another than populations with different resistance genes, suggesting that compensatory adaptation in phage-resistant bacteria also depends on evolutionary history. Several other studies have observed greater genomic parallelism among experimental evolution populations with similar starting genotypes for other traits, including in antibiotic resistance evolution (Card et al. 2021) and in compensatory adaptation after gene deletion (Rojas Echenique et al. 2019). And outside of the laboratory, convergence in sequence evolution appears to be more common among populations with a recent common ancestor than a distant one (Conte et al. 2012; Goldstein et al. 2015). Notably, the effect we observed was strong when the phage selection event was recent, but was no longer detectable after populations had spent an extended period of time in the same environment. Thus, recent historical differences appear to be more important than distant historical differences in shaping subsequent evolution.

### Conclusions

Our study provides evidence for an evolutionary ratchet in bacteria-phage coevolution, where compensatory adaptation enables the persistence of certain resistance mutations even after the original selection pressure has ceased to operate. Our findings provide important context to previous observations that phage resistance persists for long periods in some populations (Meyer et al. 2010; Wielgoss et al. 2016) but diminishes over time in others (Avrani and Lindell 2015; Wielgoss et al. 2016). In our study, the ancestral, phage-sensitive genotype was able to block phage infection through any one of at least 17 individual mutations. Given the strong selection that phages impose on their hosts, it is likely that whichever resistance mutation appears first will rapidly increase in frequency; yet the identity of this initial mutation has critical implications for the subsequent evolution and evolvability of the population.

Our findings add to a growing body of work that selection by phages can play a key role in the evolutionary dynamics of bacterial communities. Furthermore, many of the bacterial populations in our study acquired mutations in genes known to interact with eukaryotic immune systems. Phage resistance was directly mediated in many cases by changes to lipopolysaccharide molecules, which are recognized by both animal and plant immune systems (Triantafilou and Triantafilou 2005; Newman et al. 2007), and many populations also acquired mutations in type III effector proteins that underlie bacterial virulence. It will thus be important to characterize whether and how phages are indirectly responsible for shaping coevolution in between bacteria and eukaryotes as well (Wahida, Tang, and Barr 2021).

## METHODS

### Selection and validation of phage resistance

*Pseudomonas syringae* pv. tomato DC3000 was obtained from Gail Preston at Oxford. The phage strains FMS, VCM, M5.1, QAC, and SNK were obtained from OmniLytics, Inc. Phage-resistant colonies were selected through soft agar overlays. Briefly, we amplified *P. syringae* in King’s B medium overnight, then mixed bacterial cells with soft agar (King’s B medium supplemented with 0.6% agar). The mixture was spread evenly on petri dishes and allowed to dry. Droplets of high titer phage were pipetted on top. Plates were incubated for 48 hours at 28°C. Large clearing zones (plaques) appeared and expanded from the phage droplet sites, and resistant colonies were picked from plaques.

To verify resistance, each bacterial colony was streaked on hard agar plates (King’s B medium supplemented with 1.2% agar), across a high titer line of the phage strain it was isolated on (Burlage et al. 1998). Ancestral DC3000 was also streaked against all phages as a phage-sensitive negative control. Colonies were verified as phage-resistant if bacterial growth was uninterrupted at the phage line (Figure **S3**).

To ensure that colonies were entirely free of phage particles prior to experimental evolution, each colony was streaked on hard agar, and cells were sampled at the end of the streak to seed an overnight culture. The next day, the overnight culture was passed through a 0.22 μm filter. The filtrate was spotted on a mixture of ancestral DC3000 and soft agar, as described above. Plates were incubated for 48 hours at 28°C and checked for plaques. This entire process was repeated twice, at which point none of the filtrates produced phage plaques.

Of note, there was no effect of the identity of the selecting phage on either resistance mechanism or fitness of the bacteria; in fact, different phages often selected for resistance mutations in the same gene or even the exact same resistance mutation. Further, resistance to one phage typically conferred cross-resistance to all other phages in the study, suggesting that the phages in this study used the same receptor. We therefore aggregated data across selecting phages for the analyses in this study.

### Experimental evolution in the absence of phage

Phage-resistant colonies were passaged in King’s B medium in cell culture plates. Six phage-sensitive populations founded from ancestral DC3000 were passaged alongside phage-resistant colonies to serve as controls for adaptation to the lab environment, and each transfer also included a media-only negative control. Every 3 days, 75 uL of each population (approximately 10^6^ cells) were transferred to a new well with 4 mL of media. At every other transfer (every 6 days), a subset of each population was combined with a 50:50 mixture of King’s B medium and glycerol for storage at –80°C. The experiment lasted for 36 days (12 transfers), at which point there were no longer any detectable fitness differences between populations founded from phage-resistant colonies and populations founded from ancestral DC3000. According to previous work, the average doubling time of *Pseudomonas syringae* at this temperature is approximately 1.27 hours (Young, Luketina, and Marshall 1977).

### Resistance and fitness measurements

On days 12, 24, and 36 of experimental evolution, 100 uL from each culture was sampled, serially diluted, and plated on hard agar. From the dilution level at which individual colonies were visible, 96 colonies were picked from each population. Each colony was streaked against the phage strain that the population was originally isolated on. After two days of incubation at 28°C, colony phenotypes were scored as resistant (uninterrupted bacterial growth across the plate), moderate (partly interrupted bacterial growth), or sensitive (fully interrupted bacterial growth at the intersection with the phage line). In populations with colonies scored as sensitive, phage sensitivity was further validated with time series growth data (Figure **S4**).

To measure population growth rates over the course of experimental evolution, time series growth data was collected using a Molecular Devices VersaMax Microplate Reader. An overnight culture of each population was initially diluted to 0.001 OD_600_ in 200 mL King’s B medium. Colonies were randomized with respect to spatial layout, and media-only wells were included as negative controls. The plate was incubated at 28°C for 40 hours with continuous shaking, with optical density readings taken at 600 nm every 5 minutes. These readings were used to fit a logistic growth model with the R package *growthrates* and estimate the intrinsic growth rate *μ*_max_. To express these values as a proportion of wild-type fitness while controlling for technical effects, the fitted values were extracted from a model that included the day of experimental evolution, the population type (phage-resistant colony or phage-sensitive control), and the location of the well within the plate.

### Whole-genome sequencing

To identify mutations contributing to phage resistance, bacterial DNA was extracted from each phage-resistant clone using the DNeasy Blood and Tissue Kit. Concentrations were measured using a Qubit 3.0 Fluorometer, and samples were concentrated if necessary using an ethanol precipitation to obtain a minimum of 10 ng/uL of DNA per sample. DNA was sequenced at a depth of 300 Mb (estimated coverage of 45.9x) on the Illumina NextSeq 2000 platform at the Microbial Genome Sequencing Center (Pittsburgh, PA, USA). To identify and track the frequencies of mutations that arose during experimental evolution, bacterial DNA was extracted, quantified, and concentrated as described above from populations at day 6 and day 36 of the experiment. DNA was sequenced at a depth of 625 Mb (estimated coverage of 95.6x) as described above.

Paired-end reads were filtered and trimmed using Trimmomatic (Bolger, Lohse, and Usadel 2014). Reads shorter than 25 base pairs were discarded, and reads with an average quality score below 20 within a 4-base pair sliding window were discarded. Pairs of reads that both passed filtering (>95% of total reads per sample) were retained. Reads were mapped to the *Pseudomonas syringae* pv. tomato DC3000 genome (BioSample accession SAMN02604017 from the Pseudomonas Genome Database) and variants were identified using *breseq* (Deatherage and Barrick 2014), a pipeline for identifying genetic variation within microbial populations. Any sites at which the genome of the ancestral strain differed from the reference genome were removed from subsequent analyses. To avoid false positive calls from repetitive regions, mutations were filtered to exclude regions of high polymorphism (five or more mutations in a 50-base pair sliding window within a population at a single time-point).

### Analysis of genomic parallelism

To quantify the genome-wide parallelism of experimentally evolved populations, a matrix was generated of all genes with newly acquired mutations in each population (i.e. those not already fixed at day 0 of the experiment). The Sørenson-Dice similarity coefficient was calculated for each pair of populations as follows, where G_1_ represents genes with newly acquired mutations in population 1, and G_2_ represents genes with newly acquired mutations in population 2.

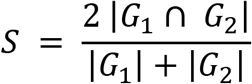

Mutations were included if they appeared at any population frequency (they were not required to be fixed), but synonymous and intergenic mutations were excluded. The analysis focused therefore on nonsynonymous point mutations or indels within coding regions.

Following (Card et al. 2021), the distribution of pairwise Sørenson-Dice values was analyzed using a randomization test. The labels annotating pairs of populations as having the same or different resistance genes were shuffled, and the mean difference was calculated between pairs with the same resistance gene and pairs with different resistance genes. This process was repeated 10,000 times to generate a null distribution. The true difference in means in the observed data was compared to the null distribution, and the observed difference was considered significant if it was more extreme than the upper 5% of permuted values.

### Replay of experimental evolution

To assess the repeatability of phage resistance outcomes in the initial evolution experiment, six populations were identified for further study. This list included three populations that re-evolved phage sensitivity in the initial experiment, each paired with the closest possible genetic match that maintained phage resistance. Ten replicates of each population were seeded from samples taken at day 0 of the initial evolution experiment (i.e. before they eventually lost or maintained phage resistance). The resulting 60 populations were maintained in King’s B medium and passaged as described above for 36 days. At this point, 96 colonies were picked from each population, individually streaked against phage, and scored as resistant or sensitive. Resistance was analyzed as a proportion of colonies within each population. Since many populations were either completely resistant or completely sensitive, Welch’s t-test for unequal variances was used to compare resistance outcomes in the replay experiment according to their founder populations.

## Acknowledgments

The authors thank members of the Koskella lab for helpful discussion and methods advice, particularly Catherine Hernandez. This work was supported by the National Science Foundation (award number 1650114 to RD; award number 1942881 to BK), the Society for the Study of Evolution (award 047408 to RD), and the Amgen Foundation (Amgen Scholars Fellowship to NDL).

## Data availability

The data and code supporting the figures in this paper can be found in the online repository https://github.com/reenadebray/loss-of-resistance.

## REFERENCES

1. Avrani, Sarit, and Debbie Lindell. 2015. “Convergent Evolution toward an Improved Growth Rate and a Reduced Resistance Range in *Prochlorococcus* strains Resistant to Phage.” Proceedings of the National Academy of Sciences 112(17): E2192–E2200. https://doi.org/10.1073/pnas.1420347112.

2. Bohannan, B. J. M., and R. E. Lenski. 2000. “Linking Genetic Change to Community Evolution: Insights from Studies of Bacteria and Bacteriophage.” Ecology Letters 3(4): 362–377. https://doi.org/10.1046/j.1461-0248.2000.00161.x.

3. Bolger, Anthony M., Marc Lohse, and Bjoern Usadel. 2014. “Trimmomatic: A Flexible Trimmer for Illumina Sequence Data.” Bioinformatics 30 (15): 2114–20.

4. Bridgham, Jamie T., Eric A. Ortlund, and Joseph W. Thornton. 2009. “An Epistatic Ratchet Constrains the Direction of Glucocorticoid Receptor Evolution.” Nature 461 (7263): 515–19.

5. Brockhurst, Michael A., Andrew D. Morgan, Andrew Fenton, and Angus Buckling. 2007. “Experimental Coevolution with Bacteria and Phage. The Pseudomonas Fluorescens--Phi2 Model System.” Infection, Genetics and Evolution: Journal of Molecular Epidemiology and Evolutionary Genetics in Infectious Diseases 7 (4): 547–52.

6. Bull, James J., Christina Skovgaard Vegge, Matthew Schmerer, Waqas Nasir Chaudhry, and Bruce R. Levin. 2014. “Phenotypic Resistance and the Dynamics of Bacterial Escape from Phage Control.” PloS One 9 (4): e94690.

7. Burlage, Robert S., Ronald Atlas, David Stahl, Gary Sayler, and Gill Geesey. 1998. Techniques in Microbial Ecology. Oxford University Press on Demand.

8. Burmeister, Alita R., Abigail Fortier, Carli Roush, Adam J. Lessing, Rose G. Bender, Roxanna Barahman, Raeven Grant, Benjamin K. Chan, and Paul E. Turner. 2020. “Pleiotropy Complicates a Trade-off between Phage Resistance and Antibiotic Resistance.” Proceedings of the National Academy of Sciences of the United States of America 117 (21): 11207–16.

9. Burmeister, Alita R., Rachel M. Sullivan, and Richard E. Lenski. 2020. “Fitness Costs and Benefits of Resistance to Phage Lambda in Experimentally Evolved Escherichia Coli*.” Evolution in Action: Past, Present and Future pp. 123–143. https://doi.org/10.1007/978-3-030-39831-6_11.

10. Card, Kyle J., Misty D. Thomas, Joseph L. Graves Jr, Jeffrey E. Barrick, and Richard E. Lenski. 2021. “Genomic Evolution of Antibiotic Resistance Is Contingent on Genetic Background Following a Long-Term Experiment with.” Proceedings of the National Academy of Sciences of the United States of America 118 (5). https://doi.org/10.1073/pnas.2016886118.

11. Clay, K., and P. X. Kover. 1996. “The Red Queen Hypothesis and Plant/pathogen Interactions.” Annual Review of Phytopathology 34: 29–50.

12. Conte, Gina L., Matthew E. Arnegard, Catherine L. Peichel, and Dolph Schluter. 2012. “The Probability of Genetic Parallelism and Convergence in Natural Populations.” Proceedings of the Royal Society B 279 (1749): 5039–47.

13. Deatherage, Daniel E., and Jeffrey E. Barrick. 2014. “Identification of Mutations in Laboratory-Evolved Microbes from next-Generation Sequencing Data Using Breseq.” Methods in Molecular Biology 1151: 165–88.

14. Dennehy, John J. 2012. “What Can Phages Tell Us about Host-Pathogen Coevolution?” International Journal of Evolutionary Biology 2012 (November): 396165.

15. Durão, Paulo, Roberto Balbontín, and Isabel Gordo. 2018. “Evolutionary Mechanisms Shaping the Maintenance of Antibiotic Resistance.” Trends in Microbiology 26 (8): 677–91.

16. Dy, Ron L., Corinna Richter, George P. C. Salmond, and Peter C. Fineran. 2014. “Remarkable Mechanisms in Microbes to Resist Phage Infections.” Annual Review of Virology 1 (1): 307–31.

17. Evans, T. J., A. Ind, E. Komitopoulou, and G. P. C. Salmond. 2010. “Phage-Selected Lipopolysaccharide Mutants of *Pectobacterium Atrosepticum* Exhibit Different Impacts on Virulence.” Journal of Applied Microbiology 109 (2): 505–14.

18. Faria, Vítor G., Nelson E. Martins, Tânia Paulo, Luís Teixeira, Élio Sucena, and Sara Magalhães. 2015. “Evolution of *Drosophila* Resistance against Different Pathogens and Infection Routes Entails No Detectable Maintenance Costs.” Evolution; International Journal of Organic Evolution 69 (11): 2799–2809.

19. Goldstein, Richard A., Stephen T. Pollard, Seena D. Shah, and David D. Pollock. 2015. “Nonadaptive Amino Acid Convergence Rates Decrease over Time.” Molecular Biology and Evolution 32 (6): 1373–81.

20. Gómez, José M., Miguel Verdú, and Francisco Perfectti. 2010. “Ecological Interactions Are Evolutionarily Conserved across the Entire Tree of Life.” Nature 465 (7300): 918–21.

21. Gong, Lizhi Ian, Marc A. Suchard, and Jesse D. Bloom. 2013. “Stability-Mediated Epistasis Constrains the Evolution of an Influenza Protein.” eLife 2 (May): e00631.

22. Gould, Stephen Jay. 1990. Wonderful Life: The Burgess Shale and the Nature of History. W. W. Norton & Company.

23. Harris, Katrina B., Kenneth M. Flynn, and Vaughn S. Cooper. 2021. “Polygenic Adaptation and Clonal Interference Enable Sustained Diversity in Experimental Pseudomonas Aeruginosa Populations.” Molecular Biology and Evolution, 38(12): 5359–5375. https://doi.org/10.1093/molbev/msab248.

24. Kircher, Martin, and Janet Kelso. 2010. “High-Throughput DNA Sequencing--Concepts and Limitations.” BioEssays: News and Reviews in Molecular, Cellular and Developmental Biology 32 (6): 524–36.

25. Koskella, Britt, and Michael A. Brockhurst. 2014. “Bacteria-Phage Coevolution as a Driver of Ecological and Evolutionary Processes in Microbial Communities.” FEMS Microbiology Reviews 38 (5): 916–31.

26. Koskella, Britt, Derek M. Lin, Angus Buckling, and John N. Thompson. 2012. “The Costs of Evolving Resistance in Heterogeneous Parasite Environments.” Proceedings of the Royal Society B 279 (1735): 1896–1903.

27. Kulikov, Eugene E., Alla K. Golomidova, Nikolai S. Prokhorov, Pavel A. Ivanov, and Andrey V. Letarov. 2019. “High-Throughput LPS Profiling as a Tool for Revealing of Bacteriophage Infection Strategies.” Scientific Reports 9 (1): 2958.

28. Lahti, David C., Norman A. Johnson, Beverly C. Ajie, Sarah P. Otto, Andrew P. Hendry, Daniel T. Blumstein, Richard G. Coss, Kathleen Donohue, and Susan A. Foster. 2009. “Relaxed Selection in the Wild.” Trends in Ecology & Evolution 24 (9): 487–96.

29. LeGault, Kristen N., Stephanie G. Hays, Angus Angermeyer, Amelia C. McKitterick, Fatema-Tuz Johura, Marzia Sultana, Tahmeed Ahmed, Munirul Alam, and Kimberley D. Seed. 2021. “Temporal Shifts in Antibiotic Resistance Elements Govern Phage-Pathogen Conflicts.” Science 373 (6554). https://doi.org/10.1126/science.abg2166.

30. Lourenço, Marta, Lorenzo Chaffringeon, Quentin Lamy-Besnier, Thierry Pédron, Pascal Campagne, Claudia Eberl, Marion Bérard, Bärbel Stecher, Laurent Debarbieux, and Luisa De Sordi. 2020. “The Spatial Heterogeneity of the Gut Limits Predation and Fosters Coexistence of Bacteria and Bacteriophages.” Cell Host & Microbe 28 (3): 390–401.e5.

31. Mangalea, Mihnea R., and Breck A. Duerkop. 2020. “Fitness Trade-Offs Resulting from Bacteriophage Resistance Potentiate Synergistic Antibacterial Strategies.” Infection and Immunity 88 (7). https://doi.org/10.1128/IAI.00926-19.

32. McCandlish, David M., Premal Shah, and Joshua B. Plotkin. 2016. “Epistasis and the Dynamics of Reversion in Molecular Evolution.” Genetics 203 (3): 1335–51.

33. McGlothlin, Joel W., Megan E. Kobiela, Chris R. Feldman, Todd A. Castoe, Shana L. Geffeney, Charles T. Hanifin, Gabriela Toledo, et al. 2016. “Historical Contingency in a Multigene Family Facilitates Adaptive Evolution of Toxin Resistance.” Current Biology 26(12): 1616–1621. https://doi.org/10.1016/j.cub.2016.04.056.

34. Meaden, Sean, Konrad Paszkiewicz, and Britt Koskella. 2015. “The Cost of Phage Resistance in a Plant Pathogenic Bacterium Is Context-dependent.” Evolution 69(5): 1321–1328. https://doi.org/10.1111/evo.12652.

35. Meyer, Justin R., Anurag A. Agrawal, Ryan T. Quick, Devin T. Dobias, Dominique Schneider, and Richard E. Lenski. 2010. “Parallel Changes in Host Resistance to Viral Infection during 45,000 Generations of Relaxed Selection.” Evolution; International Journal of Organic Evolution 64 (10): 3024–34.

36. Newman, Mari-Anne, J. Maxwell Dow, Antonio Molinaro, and Michelangelo Parrilli. 2007. “Invited Review: Priming, Induction and Modulation of Plant Defence Responses by Bacterial Lipopolysaccharides.” Journal of Endotoxin Research 13(2): 69–84. https://doi.org/10.1177/0968051907079399.

37. Pal, Csaba, María D. Maciá, Antonio Oliver, Ira Schachar, and Angus Buckling. 2007. “Coevolution with Viruses Drives the Evolution of Bacterial Mutation Rates.” Nature 450 (7172): 1079–81.

38. Pennings, Pleuni S., C. Brandon Ogbunugafor, and Ruth Hershberg. n.d. “Reversion Is Most Likely under High Mutation Supply, When Compensatory Mutations Don’t Fully Restore Fitness Costs.” bioRxiv preprint: https://doi.org/10.1101/2020.12.28.424568.

39. Picken, R. N., and I. R. Beacham. 1977. “Bacteriophage-Resistant Mutants of Escherichia Coli K12. Location of Receptors within the Lipopolysaccharide.” Journal of General Microbiology 102 (2): 305–18.

40. Rodriguez-Brito, Beltran, Linlin Li, Linda Wegley, Mike Furlan, Florent Angly, Mya Breitbart, John Buchanan, et al. 2010. “Viral and Microbial Community Dynamics in Four Aquatic Environments.” The ISME Journal 4 (6): 739–51.

41. Rojas Echenique, José I., Sergey Kryazhimskiy, Alex N. Nguyen Ba, and Michael M. Desai. 2019. “Modular Epistasis and the Compensatory Evolution of Gene Deletion Mutants.” PLoS Genetics 15 (2): e1007958.

42. Scanlan, Pauline D., Angus Buckling, and Alex R. Hall. 2015. “Experimental Evolution and Bacterial Resistance: (co)evolutionary Costs and Trade-Offs as Opportunities in Phage Therapy Research.” Bacteriophage 5 (2): e1050153.

43. Shah, Premal, David M. McCandlish, and Joshua B. Plotkin. 2015. “Contingency and Entrenchment in Protein Evolution under Purifying Selection.” Proceedings of the National Academy of Sciences of the United States of America 112 (25): E3226–35.

44. Shaver, Aaron C., Peter G. Dombrowski, Joseph Y. Sweeney, Tania Treis, Renata M. Zappala, and Paul D. Sniegowski. 2002. “Fitness Evolution and the Rise of Mutator Alleles in Experimental Escherichia Coli Populations.” Genetics 162 (2): 557–66.

45. Sheldon, B. C., and S. Verhulst. 1996. “Ecological Immunology: Costly Parasite Defences and Trade-Offs in Evolutionary Ecology.” Trends in Ecology & Evolution 11 (8): 317–21.

46. Stahl, E. A., G. Dwyer, R. Mauricio, M. Kreitman, and J. Bergelson. 1999. “Dynamics of Disease Resistance Polymorphism at the Rpm1 Locus of Arabidopsis.” Nature 400 (6745): 667–71.

47. Stenseth, Nils Chr, and J. Maynard Smith. 1984. “Coevolution in Ecosystems: Red Queen Evolution or Stasis?” Evolution 38(4): 870–880. https://doi.org/10.2307/2408397.

48. Teotónio, Henrique, and Michael R. Rose. 2007. “Perspective: Reverse Evolution.” Evolution (55(4): 653–660. https://doi.org/10.1111/j.0014-3820.2001.tb00800.x.

49. Thurber, Rebecca Vega. 2009. “Current Insights into Phage Biodiversity and Biogeography.” Current Opinion in Microbiology 12 (5): 582–87.

50. Triantafilou, Martha, and Kathy Triantafilou. 2005. “Invited Review: The Dynamics of LPS Recognition: Complex Orchestration of Multiple Receptors.” Journal of Endotoxin Research 11(1): 5–11. https://doi.org/10.1177/09680519050110010401.

51. Wahida, Adam, Fang Tang, and Jeremy J. Barr. 2021. “Rethinking Phage-Bacteria-Eukaryotic Relationships and Their Influence on Human Health.” Cell Host & Microbe 29 (5): 681–88.

52. Waterbury, J. B., and F. W. Valois. 1993. “Resistance to Co-Occurring Phages Enables Marine Synechococcus Communities to Coexist with Cyanophages Abundant in Seawater.” Applied and Environmental Microbiology 59 (10): 3393–99.

53. Weinbauer, Markus G., and Fereidoun Rassoulzadegan. 2004. “Are Viruses Driving Microbial Diversification and Diversity?” Environmental Microbiology 6 (1): 1–11.

54. Wielgoss, Sébastien, Tobias Bergmiller, Anna M. Bischofberger, and Alex R. Hall. 2016. “Adaptation to Parasites and Costs of Parasite Resistance in Mutator and Nonmutator Bacteria.” Molecular Biology and Evolution 33 (3): 770–82.

55. Woods, Robert J., Jeffrey E. Barrick, Tim F. Cooper, Utpala Shrestha, Mark R. Kauth, and Richard E. Lenski. 2011. “Second-Order Selection for Evolvability in a Large Escherichia Coli Population.” Science 331 (6023): 1433–36.

56. Young, J. M., R. C. Luketina, and A. M. Marshall. 1977. “The Effects on Temperature on Growth in Vitro of Pseudomonas Syringae and and Xanthomonas Pruni.” The Journal of Applied Bacteriology 42 (3): 345–54.

