## Supplementary Figures for "Historical contingency drives compensatory evolution and rare reversal of phage resistance"

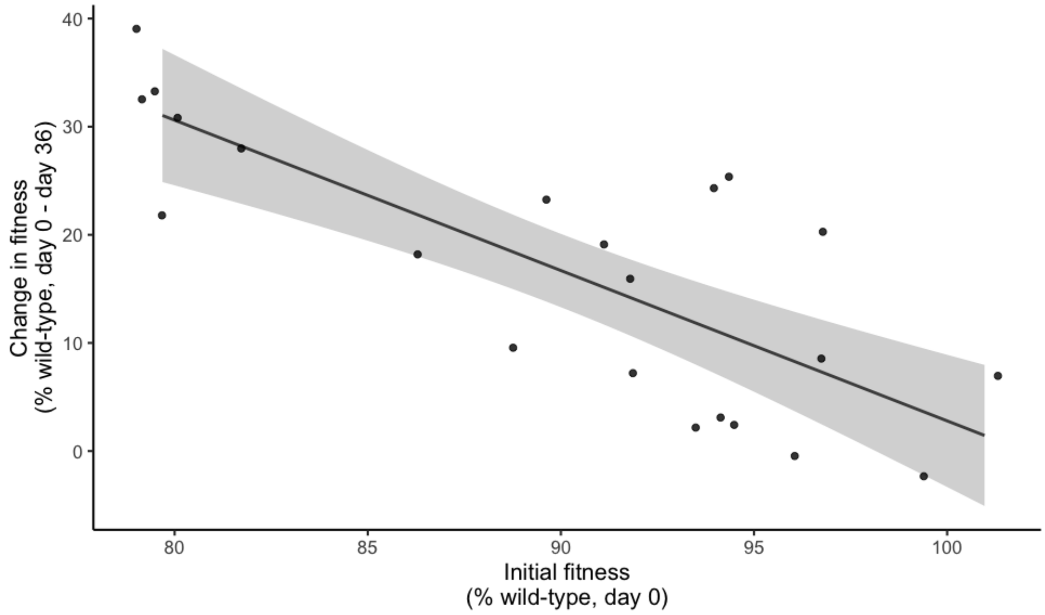


**Figure S1. Diminishing returns in adaptation over time.** Change in fitness (population growth rate) over the entire evolution experiment (days 0-36) as a function of initial fitness.

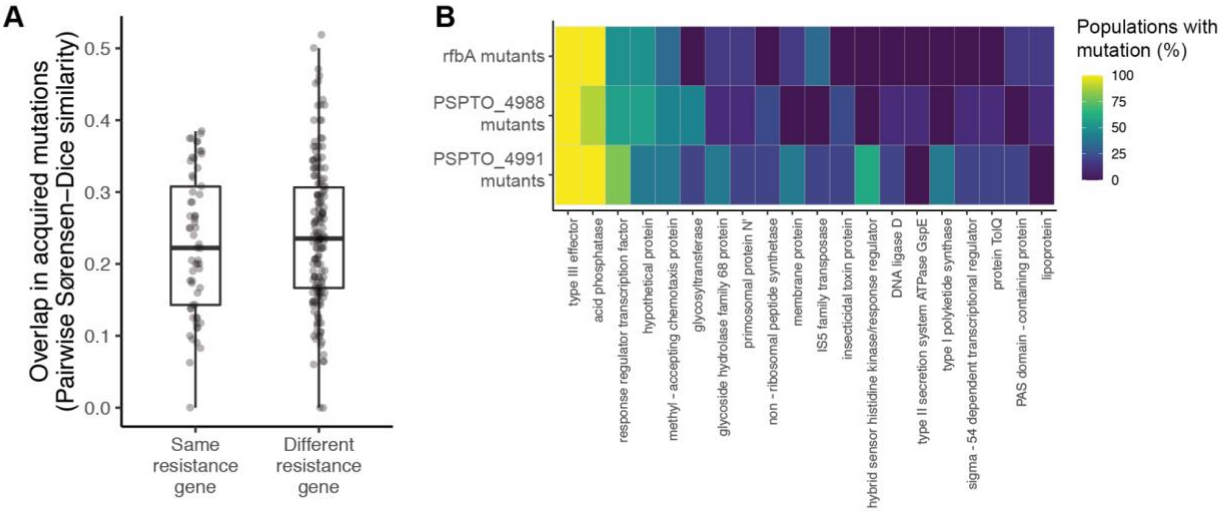


**Figure S2. Genetic contingency is no longer detectable after extended evolution in the absence of phages. (a)** Pairwise similarity coefficients among pairs of evolved populations at day 36 of experimental evolution. Statistical significance was assessed by randomizing whether pairs were labeled as having the same or different resistance genes and recalculating their similarity coefficients for 10,000 permutations. **(b)** Heatmap depicting the relationship between initial resistance genes and mutations acquired during experimental evolution. Rows represent all populations with resistance mutations in the same gene (note that resistance genes represented by fewer than 2 populations are not pictured, as there was no way to assess parallelism in these cases). Columns represent the top 20 genes that were most frequently mutated across populations by day 36 of experimental evolution. Colors indicate the percentage of populations of each resistance gene that had acquired one or more mutations by day 36 of experimental evolution.

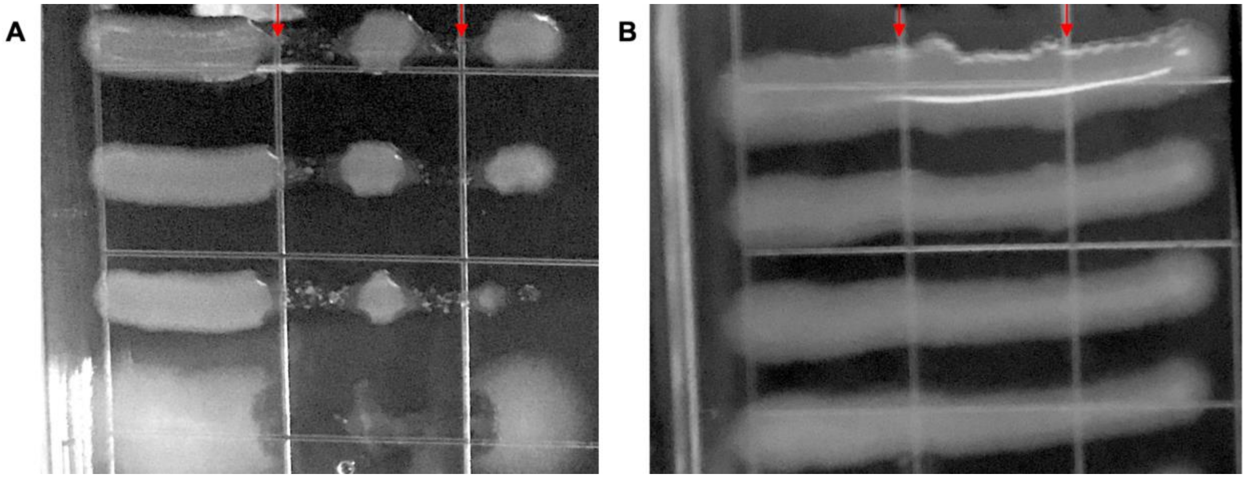


**Figure S3. Assay for phage resistance.** Two droplets of pure phage culture were pipetted at the top of each agar plate and allowed to flow downwards (red arrows). Once the droplets had dried, overnight cultures of bacterial colonies were streaked across the phage lines. Plates were incubated at 28°C for 48 hours, then colonies were scored as **(a)** phage-sensitive if bacterial growth was clearly disrupted at the phage line, or **(b)** phage-resistant if bacteria grew uninterrupted across the plate.

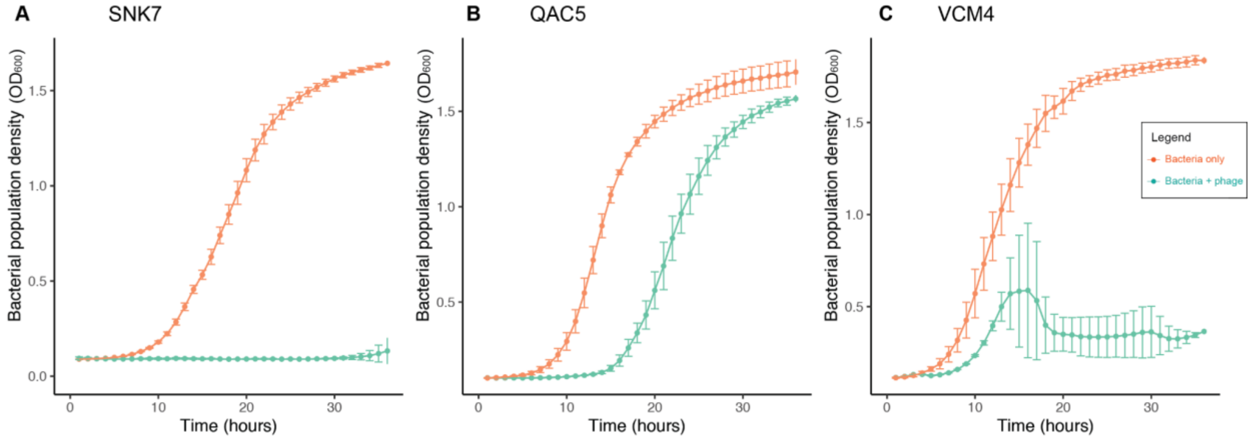


**Figure S4. Validation of phage sensitivity in evolved populations.** Growth curves of individual colonies picked from populations SNK7 **(a)**, QAC5 **(b)**, and VCM4 **(c)**. Either 30 uL of phage (overnight co-culture filtered to exclude bacterial cells) or 30 uL of spent media (overnight bacterial culture filtered to exclude bacterial cells, as a control for how phage propagation changes media composition) was added to each well. Error bars represent the standard deviation of 2-3 technical replicates per colony.
